# Intron Losses and Gains in Nematodes: Not Eccentric at All

**DOI:** 10.1101/2021.04.06.438725

**Authors:** Ming-Yue Ma, Ji Xia, Kunxian Shu, Deng-Ke Niu

## Abstract

The evolution of spliceosomal introns has been widely studied among various eukaryotic groups. Researchers nearly reached the consensuses on the pattern and the mechanisms of intron losses and gains across eukaryotes. However, according to previous studies that analyzed a few genes or genomes of nematodes, Nematoda seem to be an eccentric group. Taking advantage of the recent accumulation of sequenced genomes, we carried out an extensive analysis on the intron losses and gains using 104 nematodes genomes across all the five Clades of the phylum. Nematodes have a wide range of intron density, from less than one to more than nine per 1kbp coding sequence. The rates of intron losses and gains exhibit significant heterogeneity both across different nematode lineages and across different evolutionary stages of the same lineage. The frequency of intron losses far exceeds that of intron gains. Five pieces of evidence supporting the model of cDNA-mediated intron loss have been observed in ten *Caenorhabditis* species, the dominance of the precise intron losses, frequent loss of adjacent introns, and high-level expression of the intron-lost genes, preferential losses of short introns, and the preferential losses of introns close to 3′-ends of genes. Like studies in most eukaryotic groups, we cannot find the source sequences for the limited number of intron gains detected in the *Caenorhabditis* genomes. All the results indicate that nematodes are a typical eukaryotic group rather than an outlier in intron evolution.

## Introduction

In the nuclear genomes, protein-coding genes are often interrupted by noncoding sequences removed from the pre-mRNAs by the dynamic RNA-protein complex, spliceosome. These interrupting sequences are termed spliceosomal introns and abbreviated as introns in most publications. Eukaryotic genomes vary considerably in their intron contents. The human genome contains hundreds of thousands of introns, with each human gene has eight introns on average (Mourier and Jeffares 2003). The dinoflagellate *Symbiodinium minutum* has an even higher intron density in its genome, with up to 18.6 introns per gene (Shoguchi et al. 2013). On the other side, the yeast *Saccharomyces cerevisiae* genes have only 0.05 introns on average. The highly compacted genomes of some obligate intracellular microbes do not have any introns (Lane et al. 2007; Cuomo et al. 2012). A large-scale comparative analysis showed that the ancestors of all major eukaryotic groups and the last eukaryotic common ancestor all have intron-rich genomes, with the intron densities ranging from 53% to 74% of that in the human genome (Csuros et al. 2011). Together with this one, many studies indicate that recurrent intron losses dominated the evolution of eukaryotic genes, with a few episodes of substantial gains (Roy and Gilbert 2005a; Roy and Penny 2006b; Carmel et al. 2007; Coulombe-Huntington and Majewski 2007b; Roy and Penny 2007a; Stajich et al. 2007; Basu et al. 2008; Li et al. 2009; Worden et al. 2009; Ahmadinejad et al. 2010; van der Burgt et al. 2012; Hooks et al. 2014; Wang et al. 2014; Huff et al. 2016; Henriet et al. 2019; Lim et al. 2021).

The differential rates of intron loss and gain across eukaryotic lineages result from the differences in the rates of spontaneous mutations giving rise to new intron loss or gain events and the probability of fixing the new mutations in the genomes. Introns make it possible for one gene to code multiple proteins through alternative splicing (Gilbert 1978; Roy and Irimia 2014). Some intron sequences are recruited as regulatory elements of gene expression or harbor functional noncoding RNAs, and the splicing process might confer some benefits to the organisms by preventing some DNA damages associated transcription (Niu 2007; Wang et al. 2007; Niu and Yang 2011; Chorev and Carmel 2012; Gallegos and Rose 2015; Jo and Choi 2015; Rose 2019). Although there has been much agreement that a fraction of introns has essential biological functions, we are unsure whether the rest has any positive effects on the organisms. The most solid evidence on the beneficial impact of introns comes from the intron-poor eukaryote, *S. cerevisiae* (Parenteau et al. 2011; Bonnet et al. 2017; Morgan et al. 2019; Parenteau et al. 2019). It is very likely that the yeast genomes experienced extensive intron losses and retained only the introns that have functional roles or acquired some beneficial effects by processes like constructive neutral evolution (Doolittle et al. 2014).

On the other hand, most introns have been suggested to be slightly deleterious (Lynch 2002; Lynch and Conery 2003; Omilian et al. 2008). Thus, the fixation of the intron loss/gain events depends on natural selection efficiency, which is determined mainly by the effective population size. However, this hypothesis was not supported by the analysis of the intron gains in the genomic regions with reduced election efficiency across major eukaryotic lineages (Roy 2016). Instead, the author advocated an alternate explanation. It may be the availability of the spontaneous mutations giving rise to new introns or removing old introns that drives the evolution of intron-exon structures. The selective differences play only a minor role. Consistent with this idea, massive intron gains were observed only in genomes containing a family of transposable elements that carry splicing signals (Worden et al. 2009; van der Burgt et al. 2012; Verhelst et al. 2013; Collemare et al. 2015; Simmons et al. 2015; Huff et al. 2016; Henriet et al. 2019; Farhat et al. 2021). Meanwhile, intron loss frequency was associated with reverse transcriptase activity (Roy and Penny 2006a; Roy and Penny 2007b; Cohen et al. 2012; Zhu and Niu 2013).

The most widely cited mechanism of intron loss is via recombination of the genomic DNA with the cDNA molecules reverse-transcribed from mature mRNA (Fink 1987; Mourier and Jeffares 2003). Evidence supporting this idea, including precise intron loss, simultaneous loss of adjacent introns, preferential loss of short intron, and biased loss of introns at the 3′ side of genes have been repeatedly reported in most studied eukaryotic lineages from protists, fungi, plants to animals (Roy and Gilbert 2005b; Roy and Hartl 2006; Stajich and Dietrich 2006; Coulombe-Huntington and Majewski 2007a; Zhang et al. 2010; Yenerall et al. 2011; Hooks et al. 2014; Wang et al. 2014; Ma, Che, et al. 2015; Ma, Zhu, et al. 2015).

However, in nematodes, previous studies showed an entirely different picture of intron evolution. Phylogenetic analyses of a few genes or gene families found that the vast majority of intron changes during nematode evolution involve losses of introns individually, rather than multiple introns being lost together or gains of new introns (Robertson 1998; Cho et al. 2004). The authors advocated an alternate hypothesis. An intron could be simply lost in a mutation of genomic deletion, possibly involving nonhomologous recombination stimulated by the existence of short direct repeats at or near the two ends of an intron. Besides the individual loss of introns, this hypothesis predicts that most intron losses are not precise deletion of introns from genomic DNA but accompanied by the insertion and/or deletion (indel) of a few nucleotides into/from flanking exons. The eccentricity of nematode intron loss was further strengthened by analyzing the genome-wide alignments of *Caenorhabditis elegans* and *C. briggsae* (Kent and Zahler 2000). From the alignments between orthologous sequences of just two species, it is impossible to distinguish intron losses from intron gains. However, referring to the results of phylogenetic analysis of particular genes in nematodes, the authors believed that most of the intron changes they observed are intron losses. In total, they observed 263 changes of exact intron changes. Meanwhile, they detected 518 intron changes that cause indels to the flanking exons. That is, their results suggested that imprecise intron losses outnumbered precise intron losses in nematodes. Roy and Gilbert studied the intron losses in 684 groups of orthologous genes from seven eukaryotes, including *Homo sapiens, Drosophila melanogaster, Anopheles gambiae, C. elegans, Schizosaccharomyces pombe, Arabidopsis thaliana*, and *Plasmodium falciparum*. Evidence supporting the cDNA-mediated intron loss model, biased loss from 3′-end and adjacent intron loss, was observed. However, none of these patterns were observed in *C. elegans*, leading the authors to conclude that the intron loss process might be qualitatively different in nematodes (Roy and Gilbert 2005b). The lacking of evidence supporting the model of cDNA-mediated intron loss in nematodes was further strengthened by a later study of five *Caenorhabditis* genomes (van Schendel and Tijsterman 2013).

On the other side, studies on intron gain took an unexpected turn in nematodes. Coghlan and Wolfe (2004) compared the intron-exon structures between *C. elegans* and C. briggsae using the distantly related nematode *Brugia malayi*, two chordates (human and mouse), and two arthropods (fruit fly and mosquito) as outgroups. They found that 122 introns were recently gained introns in the two nematode genomes, and 28 of them have significant sequence identity to other introns, providing evidence for the introns′ origin. Two years later, Roy and Penny repeated the study using two newly sequenced relatives of *C. briggsae*: *C. remanei* and *Caenorhabditis* sp. 4 and showed that most of the 122 intron gains reported in one *Caenorhabditis* species are actually intron losses in other species (Roy and Penny 2006b). These results highlight the importance of the dense phylogenetic sampling of closely related species for drawing accurate inferences about intron evolution (Logsdon Jr et al. 1998).

All the previous studies on nematode intron evolution were based on a few genomes or gene families whose sequences were available at that time. With the rapid progress of genome sequencing and annotation, nearly 200 completely sequenced genomes are now available in WormBase (Howe et al. 2016; Howe et al. 2017). It is time to comprehensively revisit the nematode intron evolution based on a dense phylogenetic sampling of closely related genomes. By analyzing the introns of 104 nematode species, we carried out an extensive study on nematode intron evolution, with the molecular mechanism of intron loss in the *Caenorhabditis* branch deeply investigated.

## Results

### The Phylogenetic Tree of the Nematode Species

Using the best reciprocal basic local alignment search tool for protein (BLASTP) hits with a threshold E value of 10^−5^, we captured the orthologs that are present in over 90% of the analyzed species (104 nematode species and two outgroup species, *D. melanogaster* and *H. sapiens*), and at least in one of the two outgroup species. In total, 1577 groups of orthologous genes were obtained. The program RAxML (Randomized axelerated maximum likelihood) was used to construct the molecular consensus tree from these orthologous sequences (Stamatakis 2014). Only one node bootstrap value was 86. The others were more than 90, even most of the values (93.2%) were equal to 100. The topology structure is displayed using iTOL (Letunic and Bork 2019) (fig. 1). Each of the five major clades within the phylum Nematoda identified by Blaxter et al. (Blaxter et al. 1998) and adopted by the database WormBase (Howe et al. 2016; Howe et al. 2017) were distinctively clustered in the phylogenetic tree we constructed (fig. 1).

**Fig. 1.**
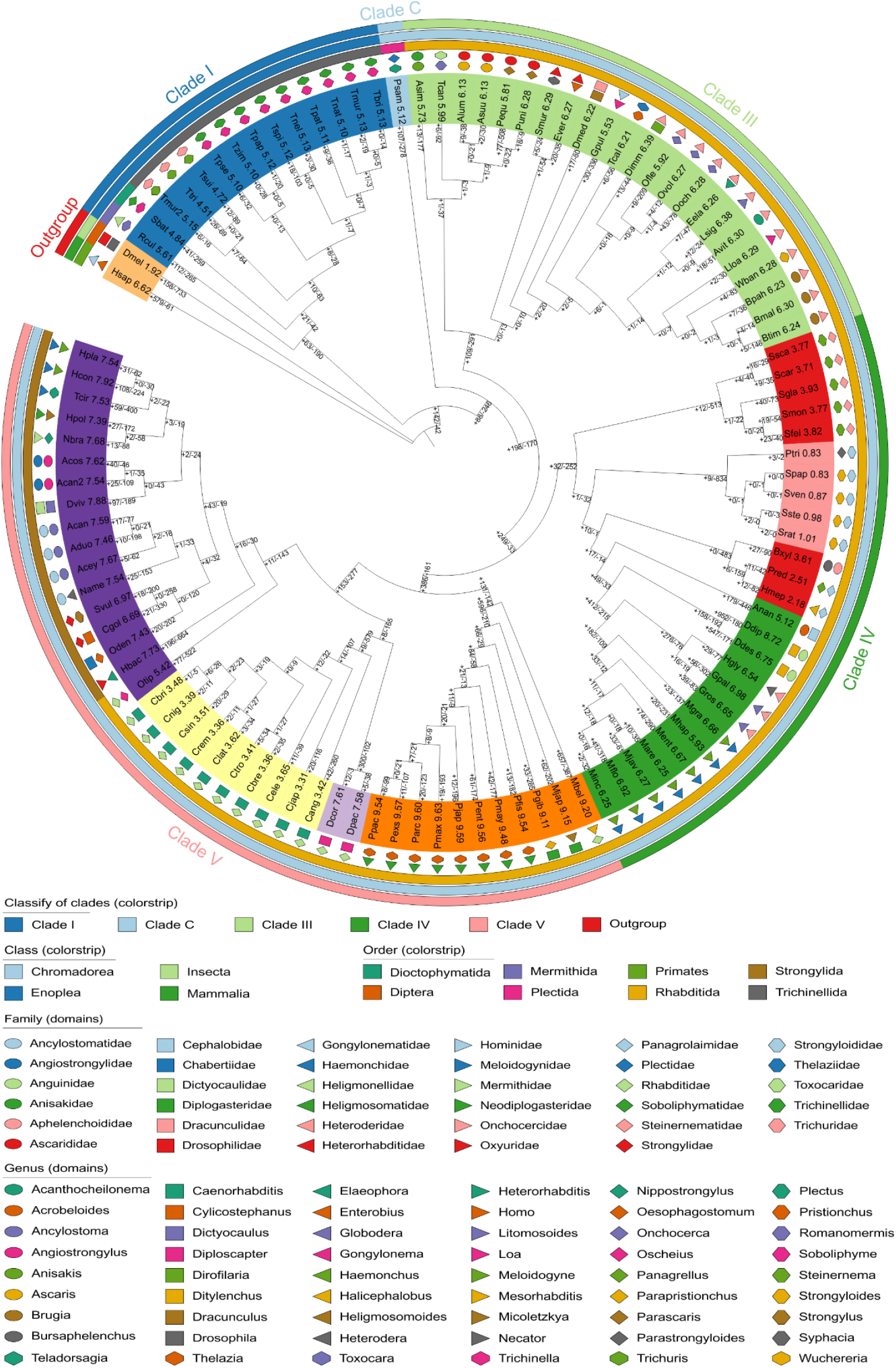
Intron losses and gains during the evolution of nematodes. This is the best tree of maximum likelihood analysis of 1577 groups of orthologous sequences. The number of intron losses and gains of each branch was computed by the Dollo parsimony method. The values are displayed on the branch lines, using “+” and “-” symbols to represent intron gain and intron loss, respectively. The numbers behind species names are intron densities. Please see Table S2 for the full name of each species and the values present in this figure. A sister figure (fig. S1) showing the rate of intron losses and gains is deposited in supplementary files.

### The Dynamic Intron Variations during Nematode Evolution

We first calculated the intron density, in the unit of the intron number per 1 kbp coding sequence (CDS), of the 1577 groups of orthologs across the 106 genomes. The intron density values of the model organisms we obtained are consistent with the previous study, with *C. elegans, D. melanogaster*, and *H. sapiens* having 3.6, 1.9, and 6.62, respectively (Csuros et al. 2011). The intron densities of the nematode genomes have a wide range, from less than one to more than nine in species (fig. 1 and Table S1). Significant differences in intron density were observed among Clade IV and Clade V species but not among Clade I or Clade III (fig. 1). The most intron-poor family Strongyloididae with intron densities ranging from 0.83 to 1.01, appears in Clade IV. At the same time, other lineages of the same clade have intron densities 2.18 to 8.72, with a median value of 6.25. The most intron-rich group, with a median value of intron density up to 9.54, is the basal taxa of the Clade V, including eight species of the family Neodiplogasteridae and three other species, *Micoletzkya japonica, Parapristionchus giblindavisi*, and *Mesorhabditis belari*. The well-studied genus *Caenorhabditis* is also presented in the Clade V. The ten *Caenorhabditis* species have a striking difference in intron density with other species in Clade V, with the median values 3.42 *vs*. 7.68. We then identified the orthologous splice sites and estimated ancestral intron content using MALIN (Csuros 2008). The number of intron loss and gain of each phylogenetic branch was calculated using the Dollo parsimony algorithm integrated into MALIN. The first pattern of nematode intron evolution that we can see is intron losses were more frequent than intron gains (fig. 1). The total number of intron losses during nematode evolution (20,270) was more than two times of intron gains (8,788). Among the 207 branches, there were 179 branches where the intron losses outnumbered intron gains. Wilcoxon Signed rank test (2-tailed) showed that the difference was highly significant (*p* = 2.3 × 10^−20^). The second pattern we observed in nematode intron evolution is that lineages with a higher intron loss rate generally have a higher intron gain rate (fig. 1 and fig. S1). Although this pattern is not so evident as the first one, statistical analysis showed is a highly significant positive correlation between the rate of intron loss and that of intron gain (Spearman’s rho = 0.68, *p* = 1.8 × 10^−29^). The third pattern of nematode intron evolution we could see from figure 1 is the vast heterogeneity in intron gain and loss rates, both across lineages and historical stages of the same lineage (fig. 1). For example, the two families Meloidogynidae and Strongyloididae, that are presented side by side in Clade IV, experienced entirely different dynamics of intron evolution. High rates of intron loss and gain constantly occurred with the lineage splitting during the evolution of the family Meloidogynidae. By contrast, massive intron losses (834) happened when the common ancestor of the family Strongyloididae branched off from other lineages. After that, very few introns were lost during line splitting events within the family. Only 9-12 intron gains happed from the common ancestor to the terminal nodes of the family Strongyloididae.

We noticed that the basal taxon, i.e., the unbranched lineages that evolved early from the root of each order or each family, generally have higher numbers of intron losses and gains (fig. 1), indicating a cumulative effect of their long evolutionary history. Therefore, we calculated the cumulative number of intron losses and that of intron gains for each species by adding the number of intron losses and gains from the root of nematodes to each tip. To avoid the effect of a common ancestor in examining their relationship, we first measured their phylogenetic signals of the current intron density of each species using the program RRR. All these three characters have very strong phylogenetic signals: λ = 0.98 (*p* = 4 × 10^−55^), 1.00 (*p* = 10^−77^), and 1.00 (*p* = 4 × 10^−59^) for the cumulative number of intron losses, the cumulative number of intron gains, and present intron density, respectively. Therefore, we performed phylogenetic generalized least squares (PGLS) for their relationships. Consistent with that obtained from individual branches, the cumulative number of intron losses is positively correlated with that of intron gains (adjusted R^2^ = 0.034, *p* = 0.033, and slope = 0.16, Table 1). Furthermore, we found that the present intron densities of the nematode species have a significant correlation with the cumulative number of intron gains (adjusted *R*^2^ = 0.4181, *p* = 7 × 10^−14^, and slope = 0.0035, Table 1), but without a statistically significant correlation with the cumulative number of intron losses (*R*^2^ = 0.025, *p* = 0.057, slope = −0.0008, Table 1).

**Table 1.** Relationships among the Frequency of Intron Losses, Frequency of Intron Gains, and Genomic Characteristics in Nematodes

| | | Slope | $R^2$ | $p$ | BH-adjusted $p$ |
| --- | --- | --- | --- | --- | --- |
| $CN_{\text{intron losses}}$ | $CN_{\text{intron gains}}$ | 0.158 | 0.035 | 0.033 | 0.096 |
| $CN_{\text{intron losses}}$ | Intron density | -0.001 | 0.025 | 0.059 | 0.127 |
| $CN_{\text{intron gains}}$ | Intron density | 0.004 | 0.418 | $7 \times 10^{-14}$ | $5 \times 10^{-13}$ |
| $CN_{\text{intron losses}}$ | Exon length | 0.005 | -0.007 | 0.592 | 0.600 |
|  | Intron length | -0.062 | 0.016 | 0.102 | 0.191 |
| | CDS size | -2.010 | 0.383 | $2 \times 10^{-12}$ | $8 \times 10^{-12}$ |
|  | Genome size | -0.107 | 0.032 | 0.038 | 0.096 |
| $CN_{\text{intron gains}}$ | Exon length | -0.062 | 0.217 | $4 \times 10^{-7}$ | $10^{-6}$ |
|  | Intron length | -0.057 | 0.003 | 0.254 | 0.381 |
|  | CDS size | 0.147 | -0.006 | 0.516 | 0.595 |
|  | Genome size | 0.036 | -0.007 | 0.600 | 0.600 |
| Intron density | Exon length | -19.04 | 0.613 | $< 2 \times 10^{-16}$ | $< 3 \times 10^{-15}$ |
|  | Intron length | 10.99 | 0.004 | 0.236 | 0.381 |
|  | CDS size | 39.49 | -0.002 | 0.386 | 0.521 |
|  | Genome size | 10.35 | -0.003 | 0.417 | 0.521 |

Note: The relationships were analyzed using phylogenetic generalized least squares analysis. The phylogenetic signals ( $\lambda$ ) are 0.98, 1.00, 1.00, 1.00, 1.00, 0.76, and 1.00 for the cumulative number of intron losses ( $CN_{\text{intron losses}}$ ), cumulative number of intron gains ( $CN_{\text{intron gains}}$ ), present intron density (intron density), median length of exon (exon length), median length of intron (intron length), median length of coding sequences (CDS size), and genome size, respectively. Except the genome size, these traits were calculated from the 1577 orthologs of each genome. $R^2$ : Adjusted *R-squared*. BH: Benjamini–Hochberg.

Furthermore, we examined whether the cumulative number of intron losses, the cumulative number of intron gains, and the current intron density are correlated with the size of introns, exons, CDSs, or genome sizes using PGLS regression analyses (Table 1). The intron evolution dynamics are not related to the intron size or genome size. Intron gains were found to have negative correlations with the exon size. We also found that the cumulative number of intron losses is negatively correlated with CDS sizes, indicating that genes encoding large proteins are less likely to lose their introns.

As multiple correlation analyses have been performed based on the same dataset, some results might be significant just by chance. Therefore, we controlled the false discovery rate using the Benjamini–Hochberg (BH) procedure and provided the adjusted *p* values in Table 1. The conclusions presented above are not changed after these corrections except the correlations of the cumulative number of intron losses with the cumulative number of intron gains and genome size.

### Intron Variations in *Caenorhabditis*

To gain insight into the mechanism of intron losses and intron gains in nematodes, we carried out an in-depth analysis of the ten *Caenorhabditis* species. In total, 4,892 groups of orthologous genes present in all the ten species were identified using BLASTP (threshold of *E* value = 10^−10^). By aligning the CDSs of orthologous genes, 6,441 discordant intron positions were detected in 2,333 groups of orthologs. Meanwhile, 6,252 conserved intron positions were identified. Some ambiguous intron positions were discarded. In 682 groups of orthologs, all the intron positions are conserved. Referring to the phylogenetic tree of the ten *Caenorhabditis* species and 12 outgroup species (fig. S2), the absence or presence of discordant introns was determined. In total, 5,047 cases of intron loss and 262 cases of putative intron gain were identified in the 10 *Caenorhabditis* species. To avoid the mis-annotations of new insertions in the transcripts into novel introns, we verified the annotation of the exon-intron structures of the 244 genes using RNA-seq data. In this way, the annotations of 168 novel introns were confirmed. Although the sample size is too small to give statistical conclusions, two evident patterns could be seen. The first is that intron loss frequency is superior to that of intron gain (Table 2). The second is a positive association between the number of intron losses and that of intron gains. The highest number of intron losses and the highest number of intron gains were detected in the basal lineage, *C. angaria*, and the lowest numbers of intron losses and intron gains were seen in recently diverged species, like *C. briggsae* (Table 2).

**Table 2.** Intron Losses and Gains in *Caenorhabditis*

| Species | Intron Losses | Intron-Lost<br>Genes | Intron<br>Gains | Intron-Gained<br>Genes | Gene with Both<br>Intron Loss and Gain |
| --- | --- | --- | --- | --- | --- |
| *C. angaria* | 2981 | 1534 | 130 | 118 | 80 |
| *C. japonica* | 918 | 672 | 16 | 15 | 7 |
| *C. elegans* | 329 | 274 | 13 | 13 | 3 |
| *C. tropicalis* | 279 | 246 | * | — | — |
| *C. brenneri* | 256 | 225 | 7 | 6 | 4 |
| *C. latens* | 41 | 38 | 0 | — | — |
| *C. remanei* | 31 | 27 | 0 | — | — |
| *C. sinica* | 166 | 142 | * | — | — |
| *C. nigoni* | 39 | 35 | 0 | — | — |
| *C. briggsae* | 7 | 6 | 2 | 2 | 0 |
| Sum | 5047 | 3199 | 168 | 154 | 94 |

Note: \*We detected six and 12 putative intron gains in *C. tropicalis* and *C. sinica*, respectively.
However, none of their annotations were confirmed by the RNA-seq data and thus all of them were not counted as novel introns.

Besides, we noticed some genes that experienced both intron losses and intron gains (Table 2). Among the 4,892 orthologous genes in *C. angaria*, the intron-lost genes (1534) takes 31.4%, and the intron-gained genes (118) take 2.41%. If intron losses and gains are randomly distributed among the genes, the genes that experienced both intron losses and intron gains should take 0.76%, i.e., 37 genes. This expected number is significantly smaller than the observed one, 80 genes Pearson’s *Chi-squared* test, *p* = 6 × 10^−18^). The same patterns were observed in *C. japonica, C. elegans*, and *C. brenneri* (Pearson’s *Chi-squared* test, *p* = 2 × 10^−4^, 0.006, and 4 × 10^−13^, respectively).

In the above analysis across 104 nematode species, we found more intron losses in species with shorter CDSs (Table 1). However, when the comparison was performed within each *Caenorhabditis* genome, a different pattern was observed. In all these ten species, the CDSs of the intron-lost genes are consistently longer than those of the intron-conserved genes (Table 3). Although the BH-correction for multiple comparisons is often suggested to be too conservative and might lead to false-negative results, all the BH-adjusted *p* values are statistically significant.

**Table 3.** Comparing the Gene Structures between the Intron-Lost Genes and Intron-Conserved Genes of *Caenorhabditis*

| Species | Gene type<sup>1</sup> | Gene number | Coding sequence length |  | Gene length |  | Intron number |  |
| --- | --- | --- | --- | --- | --- | --- | --- | --- |
| | | | Median (bp) | $p$-value<sup>2</sup> | Median (bp) | $p$-value<sup>2</sup> | Median | $p$-value<sup>2</sup> |
| Cang | Conserved | 682 | 767 | $P_U=10^{-67}$ | 1791 | $P_U=7\times10^{-30}$ | 3 | $P_U=2\times10^{-30}$ |
| | Lost | 1534 | 1350 | $P_{BH}=10^{-66}$ | 2785 | $P_{BH}=3\times10^{-29}$ | 5 | $P_{BH}=2\times10^{-29}$ |
| Cjap | Conserved | 682 | 899 | $P_U=2\times10^{-56}$ | 2860 | $P_U=6\times10^{-31}$ | 4 | $P_U=4\times10^{-24}$ |
| | Lost | 672 | 1572 | $P_{BH}=9\times10^{-56}$ | 4842 | $P_{BH}=6\times10^{-30}$ | 6 | $P_{BH}=2\times10^{-23}$ |
| Cele | Conserved | 682 | 1022 | $P_U=4\times10^{-32}$ | 2673 | $P_U=5\times10^{-15}$ | 5 | $P_U=3\times10^{-13}$ |
| | Lost | 274 | 1754 | $P_{BH}=10^{-31}$ | 4375 | $P_{BH}=10^{-14}$ | 7 | $P_{BH}=8\times10^{-13}$ |
| Ctro | Conserved | 682 | 945 | $P_U=2\times10^{-28}$ | 1586 | $P_U=10^{-14}$ | 4 | $P_U=3\times10^{-12}$ |
| | Lost | 246 | 1623 | $P_{BH}=3\times10^{-28}$ | 2480 | $P_{BH}=3\times10^{-14}$ | 6 | $P_{BH}=6\times10^{-12}$ |
| Cbre | Conserved | 682 | 987 | $P_U=3\times10^{-33}$ | 1956 | $P_U=2\times10^{-15}$ | 5 | $P_U=4\times10^{-14}$ |
| | Lost | 225 | 1797 | $P_{BH}=9\times10^{-33}$ | 3031 | $P_{BH}=8\times10^{-15}$ | 6 | $P_{BH}=10^{-13}$ |
| Clat | Conserved | 682 | 999 | $P_U=10^{-7}$ | 2098 | $P_U=9\times10^{-6}$ | 5 | $P_U=10^{-5}$ |
| | Lost | 38 | 1733 | $P_{BH}=2\times10^{-7}$ | 3742 | $P_{BH}=10^{-5}$ | 8 | $P_{BH}=2\times10^{-5}$ |
| Crem | Conserved | 682 | 1010 | $P_U=5\times10^{-5}$ | 2255 | $P_U=0.0005$ | 5 | $P_U=0.0007$ |
| | Lost | 27 | 1488 | $P_{BH}=6\times10^{-5}$ | 3695 | $P_{BH}=0.0006$ | 7 | $P_{BH}=0.0008$ |
| Csin | Conserved | 682 | 987 | $P_U=5\times10^{-22}$ | 1893 | $P_U=2\times10^{-12}$ | 5 | $P_U=3\times10^{-12}$ |
| | Lost | 142 | 1794 | $P_{BH}=8\times10^{-22}$ | 3111 | $P_{BH}=4\times10^{-12}$ | 7 | $P_{BH}=6\times10^{-12}$ |
| Cnig | Conserved | 682 | 1017 | $P_U=10^{-5}$ | 2260 | $P_U=0.0078$ | 5 | $P_U=0.0002$ |
| | Lost | 35 | 1584 | $P_{BH}=10^{-5}$ | 2889 | $P_{BH}=0.0080$ | 8 | $P_{BH}=0.0002$ |
| Cbri | Conserved | 682 | 1040 | $P_U=0.0045$ | 2714 | $P_U=0.0080$ | 5 | $P_U=0.0141$ |
| | Lost | 6 | 1880 | $P_{BH}=0.0045$ | 5108 | $P_{BH}=0.0080$ | 8 | $P_{BH}=0.0141$ |

Note. <sup>1</sup>The intron-lost genes are abbreviated as “lost” and the intron-conserved genes are abbreviated as “conserved” in this table. <sup>2</sup>The $P_U$ represents the $p$ value of Mann-Whitney $U$ test, the $P_{BH}$ represents the $p$ value by adjusted Benjamini–Hochberg procedure. Abbreviations: Cang, *C. angaria*; Cjap, *C. japonica*; Cele, *C. elegans*; Ctro, *C. tropicalis*; Cbre, *C. brenneri*; Clat, *C. latens*; Crem, *C. remanei*; Csin, *C. sinica*; Cnig, *C. nigoni*; Cbri, *C. briggsae*.

We also found that the intron-lost genes are consistently longer than the intron-conserved genes in all the ten species (Mann-Whitney *U* test, BH-adjusted *p* < 0.05 for all cases, Table 3). It should be noted that the lengths of the lost introns were not counted in calculating the size of intron-lost genes. Therefore, this is a stringent comparison to test the hypothesis that long genes are more likely to lose their introns than short genes.

For the preferential loss of introns from long genes, we propose that these genes might have more introns and thus more likely to lose some of their introns just by chance. Therefore, we compared the number of introns between the intron-lost genes and intron-conserved genes using the Mann-Whitney *U* test. As shown in Table 3, the intron-lost genes consistently have more introns than the intron-conserved genes in all the ten *Caenorhabditis* species (Mann-Whitney *U* test, BH-adjusted *p* < 0.05 for all cases). It is interesting to see that the intron-lost genes have significantly more introns than the intron-conserved genes even after losing some introns.

### The Mechanism of Intron Losses in *Caenorhabditis*

Among the 5047 cases of intron loss identified in the *Caenorhabditis* clade, 4844 cases (96%) are precise intron losses. The percentage of accurate intron losses in each genome ranges from 90.2%−100% (Table 4). In total, there are 828 pairs of adjacent intron losses. Resampling analysis showed that adjoining intron loss frequency is significantly higher than that expected by chance in all the ten *Caenorhabditis* species (*p* < 0.005 for all cases, Table 4).

**Table 4.** Frequency of precise intron losses and adjacent intron losses in *Caenorhabditis*

| Species | Intron losses | Precise losses | Adjacent pairs<sup>1</sup> | P-value<sup>2</sup> |
| --- | --- | --- | --- | --- |
| *C. angaria* | 2981 | 2866 | 643 | 0 |
| *C. japonica* | 918 | 882 | 119 | 0 |
| *C. elegans* | 329 | 314 | 26 | 0 |
| *C. tropicalis* | 279 | 269 | 14 | 0 |
| *C. brenneri* | 256 | 245 | 11 | $7 \times 10^{-5}$ |
| *C. latens* | 41 | 37 | 2 | 0.002 |
| *C. remanei* | 31 | 30 | 2 | 0.001 |
| *C. sinica* | 166 | 158 | 7 | $2 \times 10^{-4}$ |
| *C. nigoni* | 39 | 36 | 3 | $4 \times 10^{-5}$ |
| *C. briggsae* | 7 | 7 | 1 | 0.001 |

Note: <sup>1</sup> The number of adjacent pairs of intron losses.
<sup>2</sup> The probability of obtaining the observed adjacent pairs in random resampling for 100,000 times.

To study the length of the lost introns, we first examined whether the relative intron sizes are generally conserved among closely related species using nonparametric rank correlation analyses. As shown in Table 5, the intron lengths are significantly conserved between all paired *Caenorhabditis* species (*p* < 0.001 for all the cases). However, we noticed that correlation coefficients vary greatly, from 0.166 between *C. angaria* and *C. nigoni* to 0.902 between *C. latens* and *C. remanei*. Referring to the phylogenetic tree (fig. 1), there is a clear pattern that the correlation coefficients are higher between closely related species pairs than between distantly related species pairs. The length of lost introns could be well represented by the size of their orthologous introns in closely related species. But for early diverged species, like *C. angaria*, the length of lost introns could only be poorly represented by their orthologous introns in other *Caenorhabditis* species. Of course, the latter is also statistically acceptable. In this way, we compared the lengths of lost introns and conserved introns (Table 6). In most *Caenorhabditis* species (7/10), lost introns were significantly shorter than conserved introns (Mann-Whitney *U* test, *p* < 0.05 for all cases). But, in the other three species (*C. latens, C. nigoni*, and *C. briggsae*), no statistically significant differences were found (Mann-Whitney *U* test, *p* > 0.05 for all cases). Then, we performed this comparison within the intron-lost genes using Wilcoxon signed rank test by combining all the intron-lost genes of the ten species into one large sample. Here, we found that the lost introns are significantly shorter than the extant introns of the same gene (Wilcoxon signed rank test, *n* = 3199, *p* = 1.5 × 10^−43^, fig. 2A).

**Table 5.** Correlations of intron length between paired *Caenorhabditis* species

| Species |  | *Cang* | *Cjap* | *Cele* | *Ctro* | *Cbre* | *Clat* | *Crem* | *Csin* | *Cnig* |
| --- | --- | --- | --- | --- | --- | --- | --- | --- | --- | --- |
| *Cjap* | *rho* | 0.18 |  |  |  |  |  |  |  |  |
| | *p* | $10^{-46}$ | | | | | | | | |
| *Cele* | *rho* | 0.185 | 0.283 |  |  |  |  |  |  |  |
| | *p* | $2 \times 10^{-49}$ | $3 \times 10^{-115}$ | | | | | | | |
| *Ctro* | *rho* | 0.203 | 0.282 | 0.399 |  |  |  |  |  |  |
| | *p* | $2 \times 10^{-59}$ | $2 \times 10^{-114}$ | $5 \times 10^{-237}$ | | | | | | |
| *Cbre* | *rho* | 0.192 | 0.262 | 0.485 | 0.537 |  |  |  |  |  |
| | *p* | $5 \times 10^{-53}$ | $5 \times 10^{-99}$ | 0 | 0 | | | | | |
| *Clat* | *rho* | 0.205 | 0.282 | 0.471 | 0.535 | 0.567 |  |  |  |  |
| | *p* | $3 \times 10^{-60}$ | $10^{-114}$ | 0 | 0 | 0 | | | | |
| *Crem* | *rho* | 0.204 | 0.287 | 0.466 | 0.532 | 0.567 | 0.902 |  |  |  |
| | *p* | $6 \times 10^{-60}$ | $3 \times 10^{-119}$ | 0 | 0 | 0 | 0 | | | |
| *Csin* | *rho* | 0.197 | 0.276 | 0.503 | 0.503 | 0.579 | 0.602 | 0.599 |  |  |
| | *p* | $2 \times 10^{-55}$ | $2 \times 10^{-109}$ | 0 | 0 | 0 | 0 | 0 | | |
| *Cnig* | *rho* | 0.166 | 0.24 | 0.459 | 0.451 | 0.539 | 0.548 | 0.542 | 0.627 |  |
| | *p* | $10^{-39}$ | $7 \times 10^{-83}$ | 0 | 0 | 0 | 0 | 0 | 0 | |
| *Cbri* | *rho* | 0.171 | 0.231 | 0.459 | 0.455 | 0.54 | 0.552 | 0.544 | 0.621 | 0.865 |
| | *p* | $4 \times 10^{-42}$ | $3 \times 10^{-76}$ | 0 | 0 | 0 | 0 | 0 | 0 | 0 |

Note: $N=6252$ . Spearman rank correlation test was used for the correlation analyses. Abbreviations:
*Cang*, *C. angaria*; *Cjap*, *C. japonica*; *Cele*, *C. elegans*; *Ctro*, *C. tropicalis*; *Cbre*, *C. brenneri*; *Clat*, *C.*
*latens*; *Crem*, *C. remanei*; *Csin*, *C. sinica*; *Cnig*, *C. nigoni*; *Cbri*, *C. briggsae*.

**Table 6.** Comparison of the lengths between lost introns and conserved introns.

| Species | Lost introns |  | Conserved introns |  | *P*-value | *BH-adjusted p* |
| --- | --- | --- | --- | --- | --- | --- |
|  | (bp) |  | (bp) |  |  |  |
|  | Mean±SD | Median | Mean±SD | Median |  |  |
| *C. angaria* | 192±315 | 86 | 252±330 | 139 | $3 \times 10^{-44}$ | $3 \times 10^{-43}$ |
| *C. japonica* | 184±274 | 64 | 226±323 | 104 | $7 \times 10^{-15}$ | $4 \times 10^{-14}$ |
| *C. elegans* | 200±316 | 51 | 214±318 | 83 | $10^{-6}$ | $3 \times 10^{-6}$ |
| *C. tropicalis* | 151±254 | 50 | 225±548 | 54 | $7 \times 10^{-5}$ | 0.0001 |
| *C. brenneri* | 88±141 | 46 | 154±309 | 48 | $10^{-11}$ | $3 \times 10^{-11}$ |
| *C. latens* | 266±434 | 49 | 195±356 | 53 | 0.546 | 0.607 |
| *C. remanei* | 107±164 | 49 | 186±346 | 53 | 0.014 | 0.019 |
| *C. sinica* | 248±701 | 49 | 271±540 | 54 | 0.002 | 0.003 |
| *C. nigoni* | 677±1547 | 51 | 294±842 | 53 | 0.914 | 0.914 |
| *C. briggsae* | 239±199 | 211 | 247±413 | 53 | 0.362 | 0.453 |

*Note:* The number of lost introns could refer to the second columns of Table 2, and the number of conserved introns was 6,252. The length of lost intron was represented by orthologous presence intron length, and the conserved introns for comparison were represented by orthologous introns. The *P*-value was computed by Mann-Whitney *U* test. BH: Benjamini–Hochberg.

**Fig. 2.**
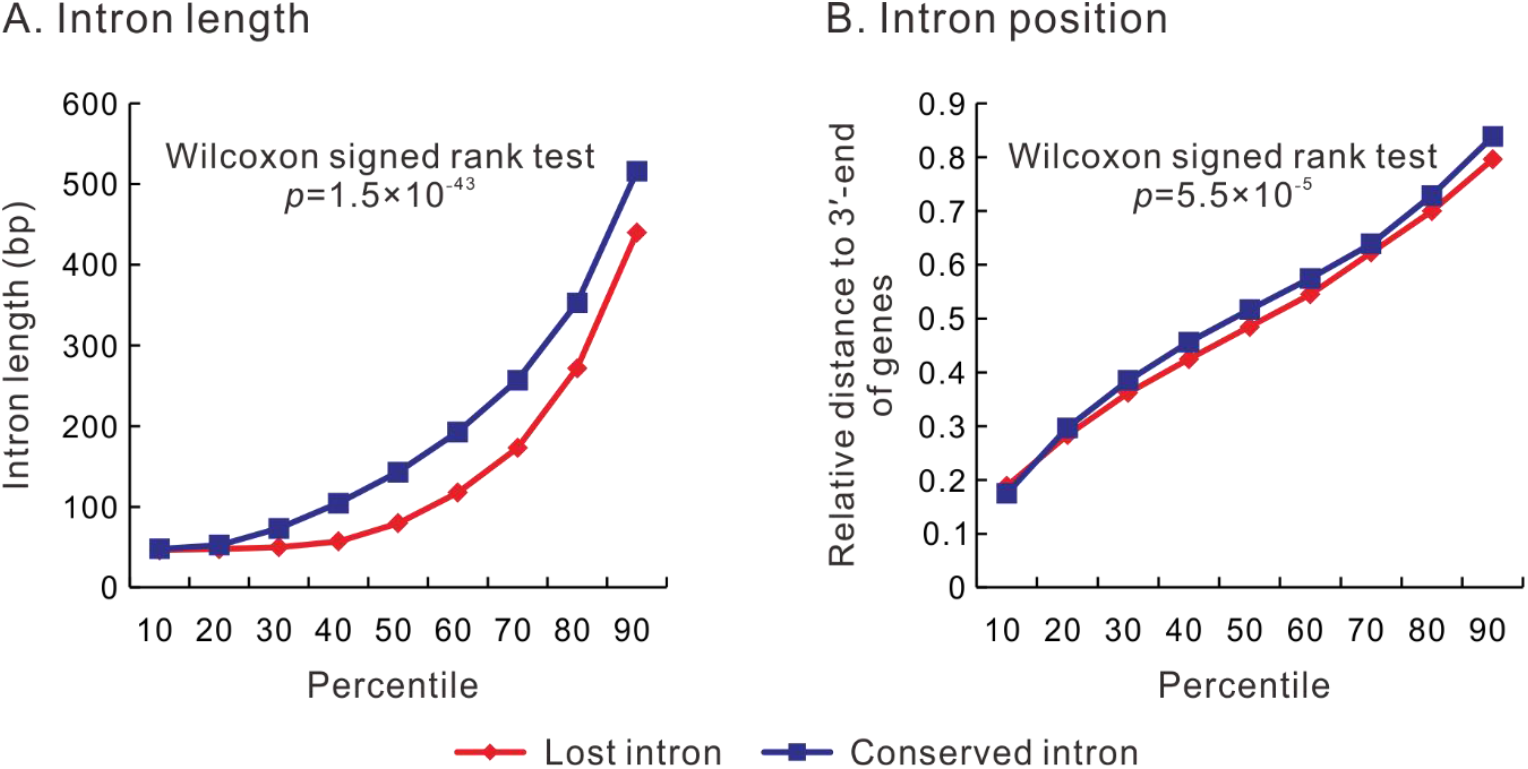
Comparison between the lost introns and conserved introns in *Caenorhabditis*. The 10th to 90th percentiles of the data are presented. (*A*) the lost introns are significantly shorter than the conserved introns of the same genes. (*B*) The lost introns are more closed to the 3′-ends of genes than conserved introns of the same genes. 3,199 intron-lost genes in *Caenorhabditis* were used in these two comparisons.

To test whether the introns close to the 3′-ends of genes were preferentially lost, we compared the lost introns and the conserved introns for their relative distance to the 3′-ends of genes which was defined as the ratio of their distances to the 3′-ends of the CDSs divided by the CDS lengths. First, we compared the lost introns with all the conserved introns in the 4,892 groups of orthologous genes using Mann-Whitney *U* tests. Only in two of the ten species (*C. japonica* and *C. remanei*), we found that the lost introns are significantly close to the 3′-ends of genes than the conserved introns (Wilcoxon signed rank test, BH-adjusted *p* < 0.05 for both cases). Then, we confined this comparison within the intron-lost genes. We averaged the relative distances of the conserved introns and those of the lost introns for each intron-lost gene. Although the mean values and median values of the lost introns are consistently smaller than those of the conserved introns, statistically significant differences were not observed in any species after the BH-correction for multiple comparisons (Wilcoxon signed rank test, *p* > 0.05 for all these three cases). However, when all the intron-lost genes from different species are considered together, they exhibit a significant difference: the lost introns are closer to the 3′-ends of genes than the current introns of the same genes, with the median values of the relative distance to the 3′-ends of genes being 0.485 and 0.517, respectively (Wilcoxon signed rank test, *p* = 5.5 × 10^−5^, fig. 2B).

Finally, we compared the expression levels of the intron-lost genes and the intron-conserved genes using the fragments per kilobase of transcript per million mapped reads (FPKM) values to estimate the gene expression level. The median FPKM values of the intron-lost genes and the intron-conserved genes are 10.26 and 5.53, respectively (fig. 3). Mann-Whitney *U* tests showed that the intron-lost genes have significantly higher expression levels than intron-conserved genes (*p* < 0.05). As an mRNA with more copies in the cytoplasm is more likely to be reverse-transcribed into cDNA, the higher expression levels of the intron-lost genes could be regarded as another piece of evidence supporting the model of cDNA-mediated intron loss.

**Fig. 3.**
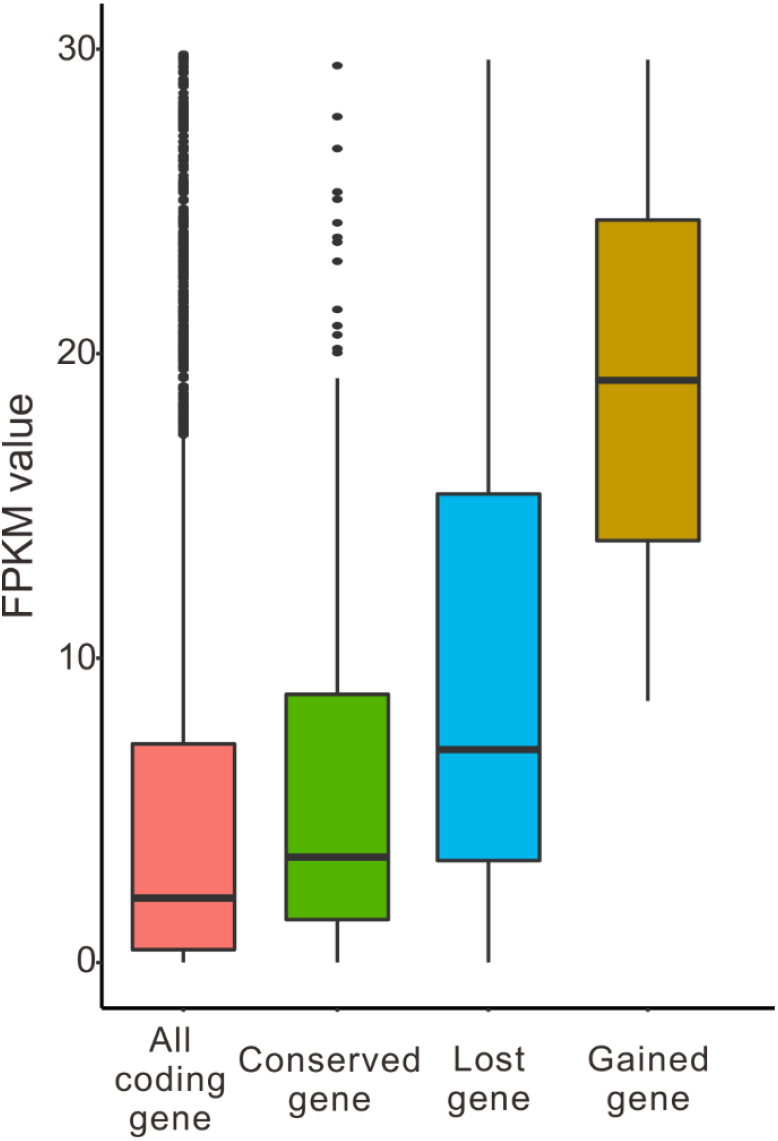
Comparison of the gene expression levels in *C. elegans*. The gene expression levels are represented by FPKM value, and the FPKM values equal to zero were removed. The numbers of all annotated coding genes, intron-conserved genes, intron-lost genes, and intron-gained genes were 3,914, 188, 50, and 2, respectively. In the box charts, the FPKM values higher than 30 are not shown. The FPKM values of the intron-lost genes are significantly higher than those of conserved genes, and of all coding genes. The *p*-values of Mann-Whitney *U* tests were 0.035 and 0.0001, respectively.

### Cases of Intron Gains in *Caenorhabditis*

From the 6,441 discordant intron positions, we identified 168 novel introns (Table S3). The newly gained introns are not evenly distributed among the ten *Caenorhabditis* species, with 130 new introns in *C. angaria*, 16 new introns in *C. japonica*, 13 new introns in *C. elegans*, seven new introns in *C. brenneri*, only two new introns in *C. briggsae*, and no new introns detected in other *Caenorhabditis* species. The 130 novel introns of *C. angaria* were distributed in 118 protein-coding genes, with five pairs of adjacent novel introns. No adjacent novel introns were found in other *Caenorhabditis* species. Hoping to find the possible source sequences for the novel introns, we searched the highly similar sequences of the novel introns in the nr/nt database and the genome database of the National Center for Biotechnology Information (NCBI). Unfortunately, no possible source sequences of the new introns were identified.

The features of the 168 newly gained introns were characterized. First, the sequence alignments flanking most (98.8%) of the freshly acquired introns did not have any gaps. These 166 introns were gained without causing insertions to or deletions from the flanking exonic sequences. Second, all the 168 new introns have the ‘gt-ag’ conservative splicing signals and the polypyrimidine tracts. One intron gain model is that the new intron was inserted into genomic DNA during DNA double-strand break repair (Li et al. 2009). Evidence supporting this model is the microhomology, or short, direct repeats flanking the gained introns, with one repeat positioned at the 5’ exon-intron boundary and the other repeat near the 3’ intron-exon boundary (Li et al. 2009). Therefore we compared the frequency of boundary-positioned microhomology between the conserved introns and the newly gained introns. Because the sample size is too small for species like *C. brenneri* and *C. briggsae*, we grouped all the 168 new introns as one sample in our comparison. The appearance of short, direct repeats was compared pairwisely between the conserved introns and the novel introns of the intron-gained genes. Short directed repeats ranging in size from three to eight base pairs were surveyed within ten bp sequences symmetrically across the exon-intron boundaries. The appearances of three to five bp short, direct repeats flanking the novel introns were significantly higher than those flanking conserved introns (Wilcoxon signed rank test, *p* = 1.4 × 10^−5^, 3 × 10^−4^, and 0.044, respectively). No significant differences were detected in the appearance of short, direct repeats ranging from six to eight bp. No statistically significant differences in GC-content between the new introns and the conserved introns were detected (Wilcoxon signed rank test, *p* = 0.57). Also, we compared the length and position of the novel introns with those of the conserved introns in intron-gained genes. The novel introns do not have significantly different sizes from the conserved introns (Wilcoxon signed rank test, *p* = 0.55) but are substantially more proximate to the 3′ ends of genes (Wilcoxon signed rank test, *p* = 0.002).

### Gene Ontology Enrichment Analysis of the Intron-Variant Genes

Taking advantage of the wealth of genomic information in *C. elegans*, we characterized the intron-lost genes and intron-gained genes for their functions and expression levels. An online gene ontology (GO) tool (http://geneontology.org/) was used to identify significant GO terms of the intron-lost genes and the intron-gained genes, with the cutoff *P*-value being set to 0.01. The significantly enriched GO terms are listed in Table S4. Most of the significant terms, biological processes, cellular components, and molecular functions, were shared by the intron-lost genes and the intron-conserved genes. Meanwhile, intron-lost genes are enriched in some particular GO terms, like ligase activity and ion binding. However, the 13 intron-gained genes do not enrich in any GO terms. Intron gains are unlikely related to specific functions.

## Discussion

With numerous studies on eukaryotic intron evolution, general patterns have been revealed. First, widespread heterogeneity in the rates of intron gain and loss have been repeatedly observed across both lineages and historical stages of the same lineage (Carmel et al. 2007; Coulombe-Huntington and Majewski 2007b; Loh et al. 2008; Farlow et al. 2010; Csuros et al. 2011; Yenerall et al. 2011). Second, intron losses were generally more frequent than intron gains, with a few episodes of burst in the intron gain rates contributed by the amplification of transposable elements carrying splicing signals (Roy et al. 2003) (Roy and Gilbert 2005a; Roy and Penny 2006b; Carmel et al. 2007; Coulombe-Huntington and Majewski 2007b; Roy and Penny 2007a; Stajich et al. 2007; Basu et al. 2008; Li et al. 2009; Worden et al. 2009; Ahmadinejad et al. 2010; van der Burgt et al. 2012; Hooks et al. 2014; Wang et al. 2014; Huff et al. 2016; Henriet et al. 2019). Third, the intron losses of most lineages are precise removals of the intron sequences from chromosomal DNA. Together with several other characteristics of intron losses, the model of the cDNA-mediated intron loss has been widely supported (Sverdlov et al. 2004; Roy and Gilbert 2005b; Stajich and Dietrich 2006; Zhang et al. 2010; Yenerall et al. 2011; Cohen et al. 2012; Zhu and Niu 2013). According to previous studies, the nematodes seem to be eccentric in their intron evolution. Their intron losses were reported to be outnumbered intron gains. Most studies failed to observe the evidence supporting the cDNA-mediated model of intron loss, like precise losses, preferential losses of adjacent introns and introns close to the 3′ end of genes, were not observed (Robertson 1998; Kent and Zahler 2000; Cho et al. 2004; Coghlan and Wolfe 2004; Roy and Gilbert 2005b; van Schendel and Tijsterman 2013).

Benefit from the unprecedented availability of genomic sequences, we carried out a large-scale, comprehensive analysis on the intron evolution of nematodes. The risk of biased observations resulting from small samples could be minimized, and a general conclusion for the intron evolution of the phylum, Nematoda, has approached. By analyzing the 104 nematode genomes, we showed that, in the aspect of intron evolution, the nematodes are a typical, rather than eccentric, group of eukaryotes. Their intron densities range from less than one to more than nine in species, almost as wide as previously reported across all eukaryotes (Mourier and Jeffares 2003). Significant heterogeneity in the rate of intron losses and gains has been observed across different nematode lineages and different evolutionary stages of the same lineage (fig. S1). Significantly more intron losses than intron gains were observed in the phylum-wide analysis and the in-depth analysis of the *Caenorhabditis* species. We examined five aspects of lost introns that are generally believed as evidence supporting the model of cDNA-mediated intron loss. In the ten *Caenorhabditis* species, the dominance of the precise intron losses, frequent loss of adjacent introns, and high-level expression of the intron-lost genes were all fully confirmed. When the lost introns and the conserved introns were compared within each species, only a few species exhibit significant differences in the preferential losses of short introns and the preferential losses of introns close to 3′-ends of genes. However, when the intron-lost genes from different species are considered together, the lost introns are significantly shorter and near the 3′-ends of genes than the extant introns. As we see, the lacking of significance in some species should be attributed to the small sample sizes. And of course, the biased position is not so strong as the other three aspects, precise intron losses, frequent loss of adjacent introns, and high-level expression of the intron-lost genes. Although the 3′-biased position was initially suggested as evidence for the cDNA-mediated intron loss (Fink 1987), it is not always observed with other evidence supporting the model (Sharpton et al. 2008; Yenerall et al. 2011). For example, in *Drosophila*, Yenerall and coauthors revealed several pieces of evidence, germline expression of the intron-lost genes, frequent loss of adjacent introns, and precise intron loss as the predominant style (Yenerall et al. 2011). However, they also found that the lost intron positions were uniformly distributed throughout the intron-lost genes. Modified versions of the cDNA-mediated intron loss that reverse transcription do not necessarily start from the 3′-end of mRNA have repeatedly been advocated in previous publications (Nielsen et al. 2004; Niu et al. 2005; Sharpton et al. 2008; Yenerall et al. 2011). The present result highlights the importance of large sample size in intron evolution studies.

The mystery of intron gains left in *Caenorhabditis* is also consistent with previous studies on other eukaryotic groups (Knowles and McLysaght 2006; Roy and Penny 2006b; Li et al. 2009; Zhang et al. 2010; Yenerall et al. 2011). No possible source sequences have been identified for the 168 novel introns detected in *Caenorhabditis*. The source sequences are regarded as the molecular smoking gun in identifying novel introns (Logsdon Jr et al. 1998; Logsdon 2004). Although researchers failed to identify the source sequences of most novel introns, they could quickly identify orthologous introns by sequence similarity (Zhang et al. 2010). As the novel introns should be gained after the divergence of the orthologous introns, there are several possible explanations for the failure in identifying source sequences. The first is that the newly acquired introns diverge at an unexpectedly high rate from their source sequences. But there is no evidence for the rapid divergence of recently gained introns in any eukaryotes. The second explanation is that the source sequences are in the dark matter that has not been sequenced. The discovery of introner elements as the novel intron sources is an example of finding a smoking gun from dark matter (van der Burgt et al. 2012; Verhelst et al. 2013; Simmons et al. 2015). Meanwhile, an insight that could be learned from the studies of introner-elements contributed novel introns is that the newly gained introns do not have an unexpectedly high divergent rate that makes them rapidly unrecognizable from their source sequences. It is also possible that the small number of intron gains, as compared with a large number of intron losses, are resulting from the imperfectness of the methods used in distinguishing intron losses and gains. A minor technical error ratio might shift a pattern of exclusively intron losses to the observation of predominant intron losses with a few cases of intron gains.

## Materials and Methods

We downloaded the genome sequences and annotation files of *D. melanogaster, H. sapiens*, and the 104 Nematoda species from Ensembl Metazoa 48, Ensembl 101 (http://www.ensembl.org/), and WormBase (release WBPS14, https://parasite.wormbase.org/index.html), respectively (Howe et al. 2016; Howe et al. 2017; Howe et al. 2020; Yates et al. 2020). The accession numbers and genomic features of the species used in this study are shown in Supplementary Table S1.

### The Orthologous Genes

The orthologous genes of the 106 species were identified using BLAST v2.2.26 (using parameter blastall -p blastp -F F -e 1e-5 -m 8) (Kent 2002). For regions that have multiple hits, only the two-way best reciprocal BLAST hits were retained. Besides, only the orthologs present in over 90% (96/106) of species were used in the subsequent analyses. In total, 1,577 sets of orthologs were obtained.

The ten *Caenorhabditis* species were selected to analyze intron evolution mechanisms and models, including *C. angaria, C. brenneri, C. briggsae, C. elegans, C. japonica, C. latens, C. nigoni, C. remanei, C. sinica*, and *C. tropicalis*. The orthologs of *Caenorhabditis* were also identified using the two-way best reciprocal BLAST hits (E value threshold = 10^−10^) (Kent 2002). A total of 4,892 sets of one-to-one orthologous genes were identified.

### The Phylogenetic Tree

The protein sequences were aligned using CLUSTALW (version 2.1) (Larkin et al. 2007). The Gblocks program (version 0.91) was used to eliminate poorly aligned regions (Castresana 2000). The filtered coding alignments were used to build the phylogenetic tree using RAxML (version 8.2.12) (Stamatakis 2014), using parameters -f a -x 1533 -# 1000 -m GTRGAMMAX -s sequences.phy -q partitions.txt.

### Inference of Ancestral Introns

We inferred the ancestral introns from 1,577 sets of orthologs of the 104 nematode genomes using the MALIN package (Csuros 2008). Firstly, we generated a table of intron presence or absence in the orthologs using MALIN. It included 10,469 intron sites allowing a maximum of 11 ambiguous entries per site. Intron loss and gain rates were optimized in MALIN using maximum likelihood with a constant rate with default parameters and running 1,000 optimization rounds (likelihood convergence threshold = 0.001). The ancestral intron presence, the numbers of intron losses and gains were estimated in MALIN using the Dollo parsimony method (Supplementary Table S2).

### Intron Variation Analysis in *Caenorhabditis*

The coding sequences of orthologs of *Caenorhabditis* species were aligned using CLUSTALW (Larkin et al. 2007) and MUSCLE (version 3.8.31) (Edgar 2004). The intron presence/absence state of orthologous alignment was compared using Perl. Only when the introns present in all the ten species were designated as a conserved intron position (6,252 conserved intron positions). The candidate intron discordant positions must meet three constraints. Firstly, the gaps within 45 bp alignment sequences upstream and downstream the intron variation positions were both less than 10. Secondly, identities of the 45 bp alignment upstream and downstream the intron variation positions were both more than 0.5. Lastly, the intron-variation-genes with at least one conserved intron position were retained. In total, 6,441 cases of discordant introns were detected.

Intron loss and potential intron gain were identified by Dollo and polymorphism parsimony algorithm (version 3.697), using parameter parsimony method = Dollo, and the input tree was shown in figure S2.

### Resampling Analysis for the Simultaneous Loss of Adjacent Introns

The probability of simultaneous intron losses was estimated using the random sampling principle. For example, in *C. elegans*, there were 26 pairs of adjacent intron-lost sites among 329 cases of intron loss. The 36,569 extant intron positions of coding sequence of 4,892 orthologous genes and 329 intron-lost positions were used to build sampled population intron positions database. Then we randomly resampled 329 positions from the 36,898 intron positions for 100,000 times. The null hypothesis is that, intron loss was a random event and so the pair-number of obtained adjacent positions should be, on average, close to 26. On the contrary, if the pair-number was much fewer than 26 in most cases, adjacent introns tends to be lost more frequently than randomly. The probability (*p*-value) was estimated by calculating the sampling times with more adjacent positions (pairs ≥ 26) divided by the total sampling times (100,000). In 100,000 times random sampling, no one sampling result of adjacent pairs was higher than or equal to 26. As a result, the probability was zero divided by 100,000. It refused the null hypothesis, and introns at different positions was not lost randomly.

### The Representative Lengths of Introns

For a lost intron, its length was represented by the length of its orthologous intron in the most closely related species. The phylogenetic relationships refer to figure S2. For instance, the intron at the orthologous position of *C. remanei* was taken as a representative intron of the lost intron of *C. latens*. For the introns in *C. sinica*, the introns at orthologous positions in *C. briggsae* and *C. nigoni* were considered as representative introns. The representative length was the average length of the orthologous introns in *C. briggsae* and *C. nigoni*. In the length comparison between lost introns and conserved introns, representative lengths were also used for conserved introns.

### Microhomology Identification

Microhomology is defined as a pair of short, direct repeats around each end of the recently gained introns. Ten bp sequences symmetrically across the exon-intron boundaries of targeted introns between upstream and downstream were surveyed for the presence/absence of microhomology. We sequentially extracted the sequences of set repeat size from the upstream and downstream boundary-positioned sequences and compared the similarities between the two regions. Only the two sequences with entirely consistence were regarded as homologous repeats. The repeat sizes were set from 3 to 8.

### Analysis of RNA-Seq Data

With abundant food, optimal temperature (20°C), and sparse population, worm development from embryo to adult, can be divided into four larval stages, L1 to L4 (Altun et al. 2002-2021). SRR7781209 and SRR7781210 were obtained at the L4-early adult stage of *C. elegans*. RNA-seq reads from *C. briggsae* (SRR7781208), *C. remanei* (SRR7781207, SRR7781212), and *C. brenneri* (SRR7781211) also obtained at the L4-early adult stage. The RNA-seq files were downloaded from the NCBI SRA database (https://www.ncbi.nlm.nih.gov/sra). A list of RNA-seq datasets used is available in Supplementary Table S5.

RNA-Seq reads were aligned to the reference genomes using TopHat algorithm v2.0.14 (using parameters --library-type fr-unstranded --min-segment-intron 10 --max-segment-intron 20000) (Kim et al. 2013). The mapped reads were used to reannotate the exon-intron structure.

The count data of RNA-seq were normalized to Reads Per Kilobase per Million (RPKM) mapped reads using Cufflinks v2.2.1, an open-source software program, using parameters -G --library-type fr-unstranded (Trapnell et al. 2013).

### Statistical Analysis

Data calculations were performed using Perl programming language. Statistical analysis and plotting were performed using R v4.0.3 and SPSS R26.0.0. Chi-square test (chisq.test function), Mann-Whitney *U* test (wilcox.test), Benjamini–Hochberg test (p.adjust function), phylogenetic signals, and PGLS were calculated using the R system, phytools v0.7-70 (Revell 2012), ape v5.4-1 (Paradis et al. 2004), MASS v7.3-53 (https://www.rdocumentation.org/packages/MASS/versions/7.3-53), mvtnorm v1.1-1 (Genz and Bretz 2009), and caper v1.0.1(Orme et al. 2011). The phylogenetic signals of seven features (the cumulative number of intron losses, the cumulative number of intron gains, current intron density, exon length, intron length, coding sequences length, and genome size) were examined using phylosig functions (using parameter method = lambda) in the R package phytools v0.7-70. Spearman rank correlation test and Wilcoxon signed ranks test were calculated using the SPSS.

## Supporting information

All the supplementary tables and figures.

## Supplementary Material

Supplementary data are submitted along with the main text.

## Acknowledgments

This work was supported by the National Natural Science Foundation of China [grant numbers 31701093, 61872115, 31671321) and the Chongqing Research Program of Basic Research and Frontier Technology (grant number cstc2017jcyjAX0200).

## Author Contributions

MYM, KS, and DKN conceived the study. MYM and JX performed the data analysis. MYM and DKN wrote the manuscript. All authors read, improved, and approved the final manuscript.

