## Supplementary material for "Intron Losses and Gains in Nematodes: Not Eccentric at All": All the supplementary tables and figures.: Supplemental figures.docx

Ming-Yue Ma^1^, Ji Xia^1^, Kun-Xian Shu^*,1^, Deng-Ke Niu^*,2^

^1^ Chongqing Key Laboratory of Big Data for Bio Intelligence, School of Bioinformatics, Chongqing University of Posts and Telecommunications, Chongqing 400065, China

^2^ MOE Key Laboratory for Biodiversity Science and Ecological Engineering and Beijing Key Laboratory of Gene Resource and Molecular Development, College of Life Sciences, Beijing Normal University, Beijing 100875, China

Fig. S2. Phylogenetic tree of *Caenorhabditis* and outgroups. The phylogenetic tree among ten *Caenorhabditis* and selected outgroup species, including the Clade I, Clade III, Clade IV, and Clade V of the Nematoda phylum, and other four metazoan species (*Rhodnius prolixus*, *Drosophila melanogaster*, *Daphnia magna*, *Crassostrea gigas*)

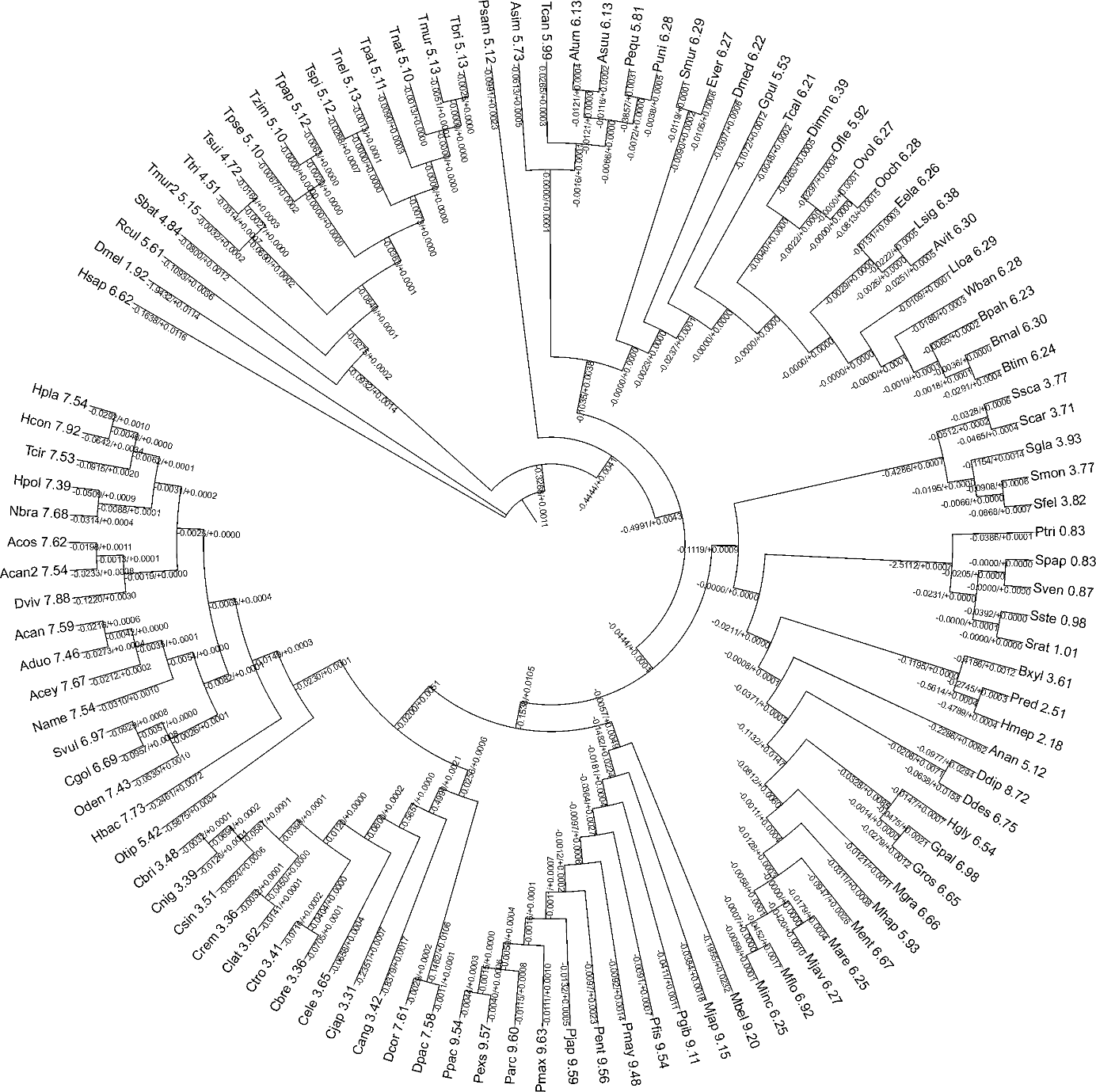

Fig. S1. The rates of intron losses and gains during the evolution of nematodes. This is the best tree of maximum likelihood analysis of 1577 groups of orthologous sequences. The rates of intron losses and gains of each branch was computed by the Dollo parsimony method. The values are displayed on the branch lines, using "+" and "-" symbols to represent intron gain rate and intron loss rate, respectively. The numbers behind species names are intron densities. Please see Table S2 for the full name of each species and the values present in this figure.

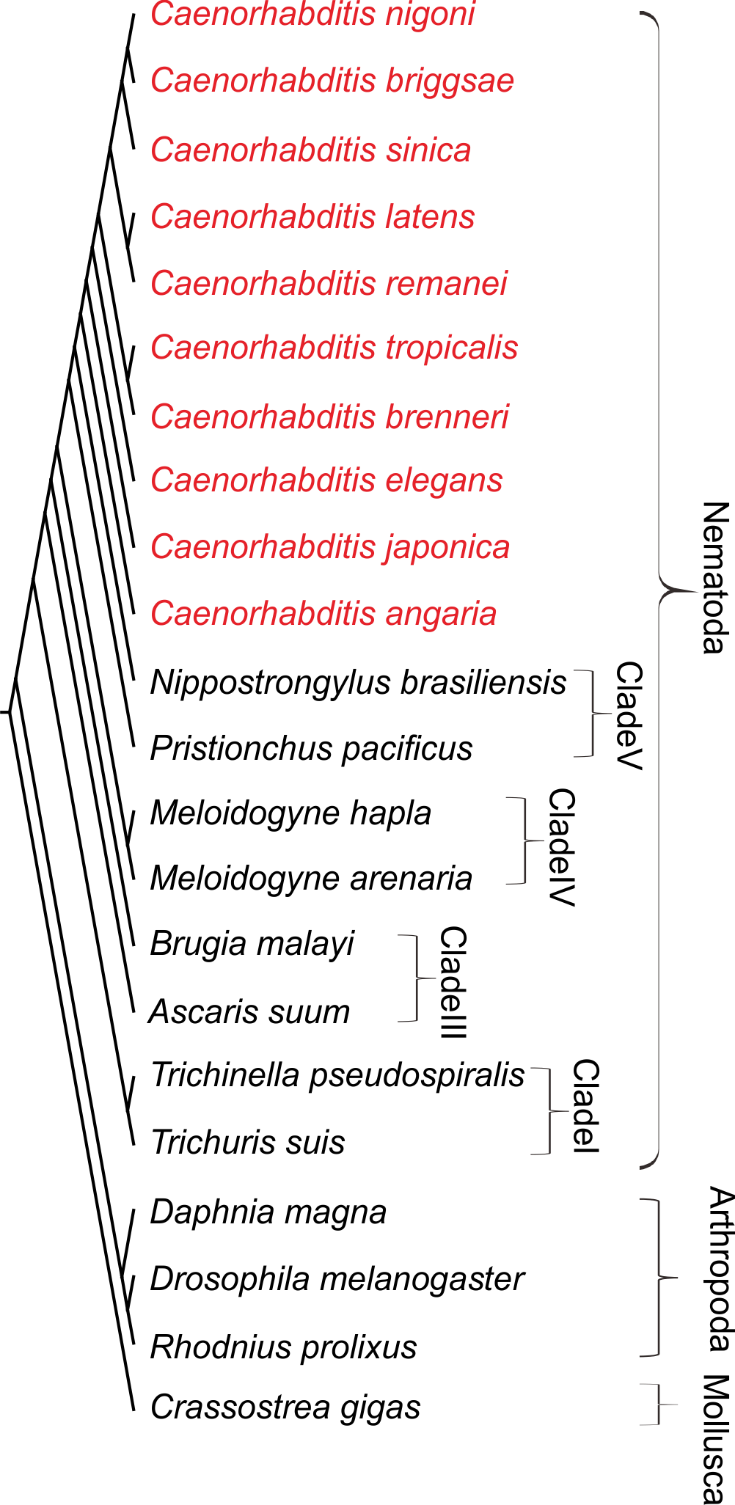
